# Distinct Temporal Patterns of Human Neural Firing in the Subthalamic Nucleus During Speech and Orofacial Movement

**DOI:** 10.64898/2025.12.11.693637

**Authors:** Zahra Jourahmad, Christopher K Kovach, Andrea H. Rohl, Joel I Berger, Kris Tjaden, Jeremy D.W. Greenlee

## Abstract

Clinical studies, along with electrophysiological findings, provide evidence that the subthalamic nucleus (STN) contributes to speech production. These studies have reported that the STN encodes diverse aspects of speech, comprising speech motor planning and execution, timing, and linguistic features such as phonetic content. However, none of these studies have included an orofacial non-speech motor task to evaluate speech-specificity of STN activity. Here, we examined the modulation of STN neurons while participants engaged in two speech tasks (sentence repetition and syllable repetition) as well as two non-speech orofacial movement tasks (jaw movement and tongue protrusion) in awake patients with Parkinson’s disease undergoing deep brain stimulation implantation surgery. A total of 51 single- and multi-unit neural clusters were captured. A Poisson generalized linear model (GLM) was implemented to understand the temporal dynamics of STN activity. A larger proportion of clusters was modulated during speech (22%) than during orofacial movement (12%) and a substantial subset of STN neural clusters responded to overlapping speech and orofacial tasks (27%). The findings suggest that STN can encode both motor and linguistic aspects of speech production.

**Graphical abstract:** The subthalamic nucleus (STN) shows neural modulation during speech production, a process which requires motor planning, execution, and phonological functions. By comparing STN spiking activity during speech tasks (sentence and syllable repetition) and non-speech orofacial tasks (jaw movement and tongue protrusion), we identified task-specific modulation patterns in STN neurons. The STN contains distinct neural populations engaged during speech and orofacial movements. Among all recorded clusters, 22% responded exclusively to speech, 12% exclusively to orofacial movements, 27% to both, and 39% were non-responsive. We demonstrated that STN activity at single- and multi-unit levels is specific to speech production and is influenced by task-specific motor and linguistic demands, highlighting a role for STN in integrating motor control and speech production.

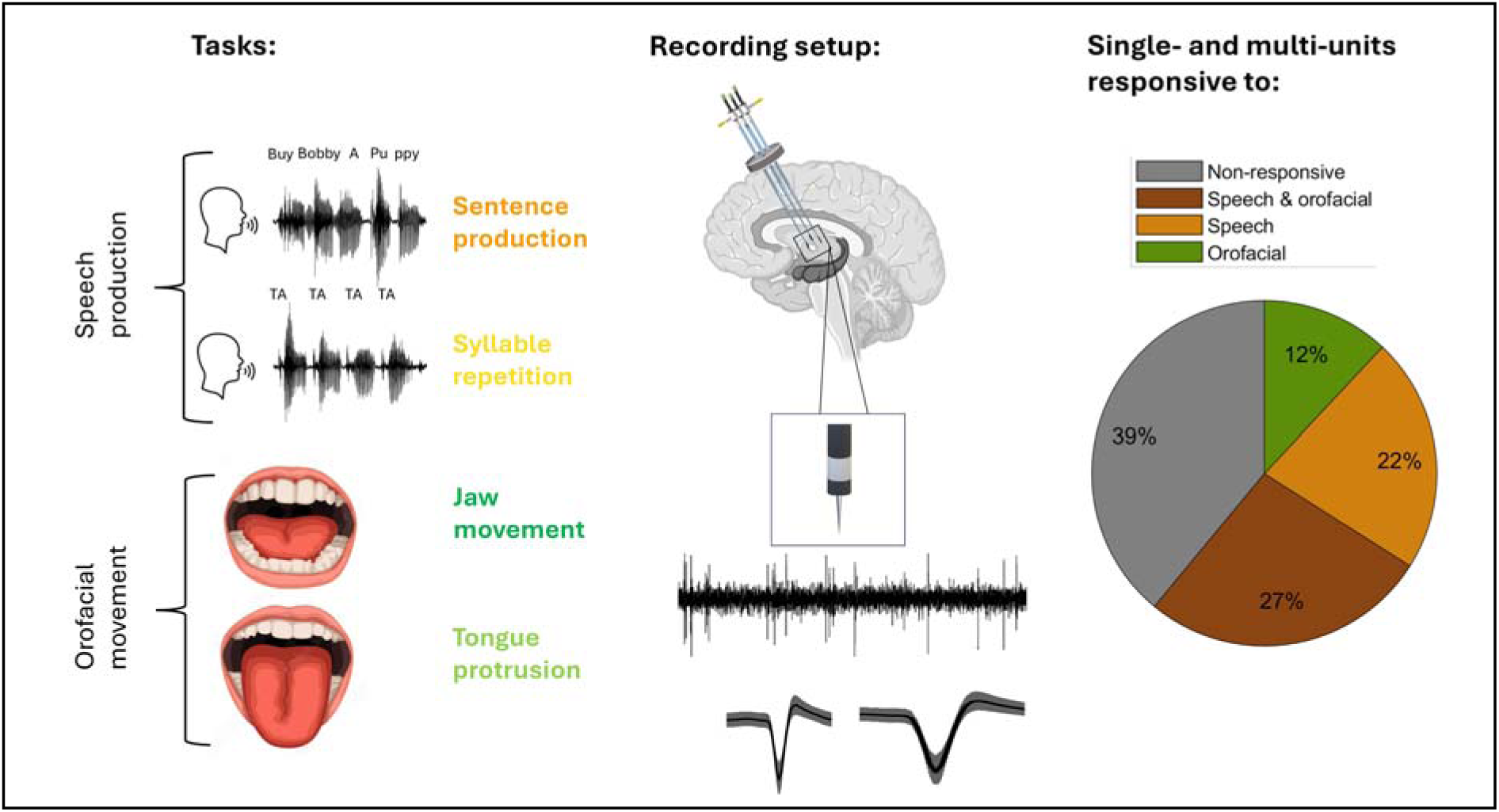

## Introduction

Speech production is a complex human behavior requiring the coordination of motor and linguistic functions. The interaction of various cortical brain regions throughout speech production is broadly studied; however, research investigating the role of subcortical areas in speech production is more limited (Friederici, 2011). Yet, speech and communication difficulties are common in disorders associated with the disruption of basal ganglia structures. For example, the manifestation of speech impairments in Parkinson’s disease (PD) is common, significant and sometimes present even in the early stages of the disease (Rohl et al., 2022). PD-related speech impairments include hypophonia, dysarthria and reduced speech intelligibility (Tabari et al., 2024; Tjaden, 2008).

Direct recordings from human subthalamic nucleus (STN) have demonstrated changes in STN firing during speech production(Lipski et al., 2018,; Lipski et al., 2024;Tankus & Fried, 2019a; Watson & Montgomery, 2006). These recordings leverage research opportunities present in awake surgery to implant deep brain stimulation (DBS) electrodes into STN for PD treatment. Previous studies employing these direct brain recordings have uncovered connections between the STN and cortical brain regions involved in speech production (Chrabaszcz et al., 2019), as well as speech perception and planning (Jorge et al., 2021) via local field potential (LFP) recordings. Moreover, single- and multi-unit neural recordings have revealed changes in the spiking activity of STN neurons during speech tasks. These studies have explored various facets of speech production, such as motor planning (Lipski et al., 2018), motor execution (Lipski et al., 2018), phonetic content(Lipski et al., 2024; Tankus & Fried, 2019a), and speech timing (Lipski et al., 2018; Watson & Montgomery, 2006).

Modulation of STN function via DBS is also known to affect speech production, representing another line of evidence for a role of STN. Speech outcomes after STN-DBS are variable and unpredictable ranging from worsening or no improvement in dysarthria to mixed outcomes in intelligibility and articulatory accuracy (Aldridge et al., 2016; Deuschl et al., 2006; Guenther & Vladusich, 2012; Müller et al., 2025; Rohl et al., 2022) despite consistent and dramatic improvements in non-speech motor symptoms after STN-DBS (Benabid et al., 2009). These clinical observations suggest that STN plays a role in producing speech. In addition, neuroimaging studies including PET and fMRI support a role for STN in speech production (Atkinson-Clement et al., 2017; Janssen & Mendieta, 2020; Manes et al., 2014; Narayana et al., 2009).

Despite the extant data outlined above, STN is not robustly included in most neural models of speech production. Such models provide frameworks for the cognitive, sensory and motor processes involved in turning thoughts into articulated speech by outlining cortico-cortical and cortico-subcortical networks (Nozari, 2022). Among models of speech production, the DIVA model includes multiple subcortical structures in shaping the model’s understanding of speech motor control and the coordination of articulatory gestures in producing phonetic content (Guenther, 2016). While the basal ganglia is recognized as a crucial subcortical structure in DIVA’s model of speech production, it is noteworthy that STN is not specifically incorporated into this model (Manes et al., 2024).

Importantly, no studies of STN physiology to date have included an appropriate control task, namely non-speech orofacial movement to mimic what occurs during speech production, to help elucidate whether the role of STN in speech production is speech-specific or reflecting orofacial motor processes. In the present study, we addressed this knowledge gap by recording STN neuronal activity at single and multi-unit levels in 12 patients with PD undergoing STN-DBS implantation surgery. We used speech and non-speech orofacial tasks (e.g., tongue, jaw, and mouth movements without phonation) to test the hypothesis that STN modulation during speech production is indeed speech-specific. Additionally, we explored whether clinical measures such as PD severity or duration of disease contribute to modulation of neural firing rates in STN for speech and orofacial movement.

## Materials and methods

### Participants

Participants were 12 patients (3 female; mean age 64 ± 6 years) with PD who were undergoing awake bilateral STN-DBS stereotactic lead implantation surgery (**Table 1**). Presurgical evaluation of participants included UPDRS-III testing to assess the severity of PD, neuropsychological assessment to exclude dementia, levodopa challenge, and structural brain MRI. None of the participants took their usual PD medications for 8-12 hours before the intraoperative experiment. The University of Iowa’s Institutional Review Board (IRB) approved this study, and all participants provided written informed consent prior to research participation. In accordance with the IRB protocol, the overall experiment duration lasted less than 15 minutes.

**Table 1.**
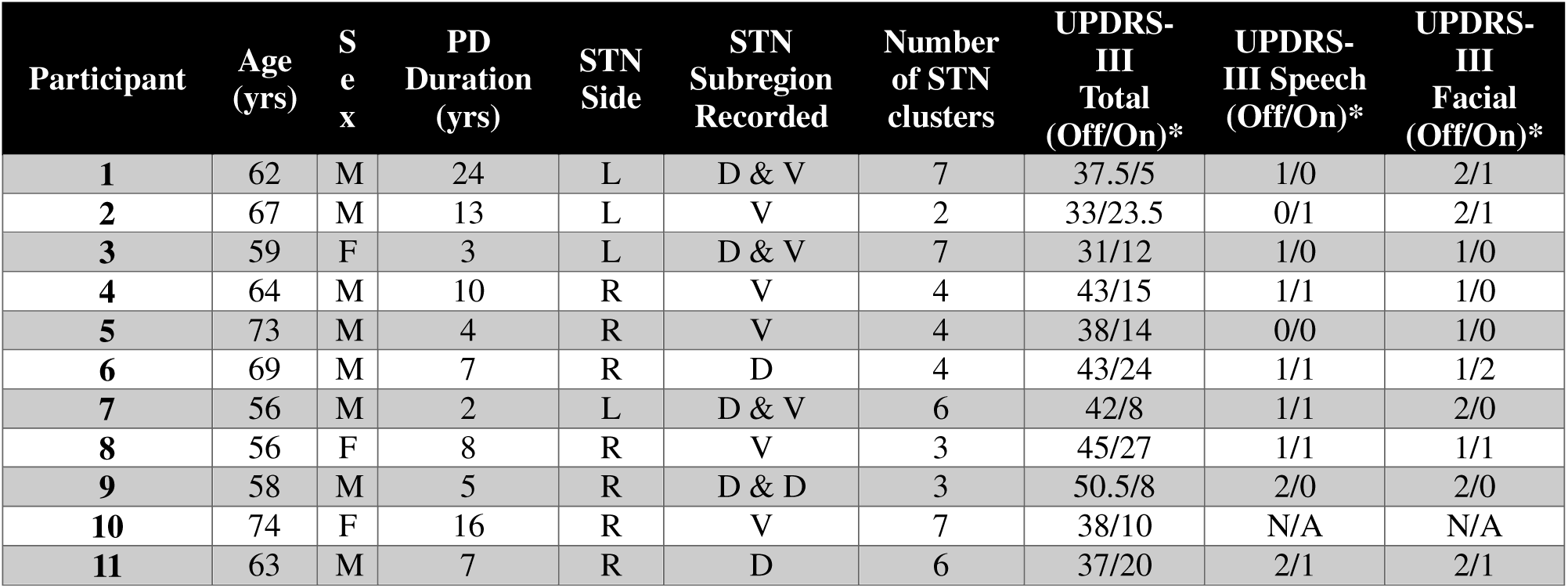
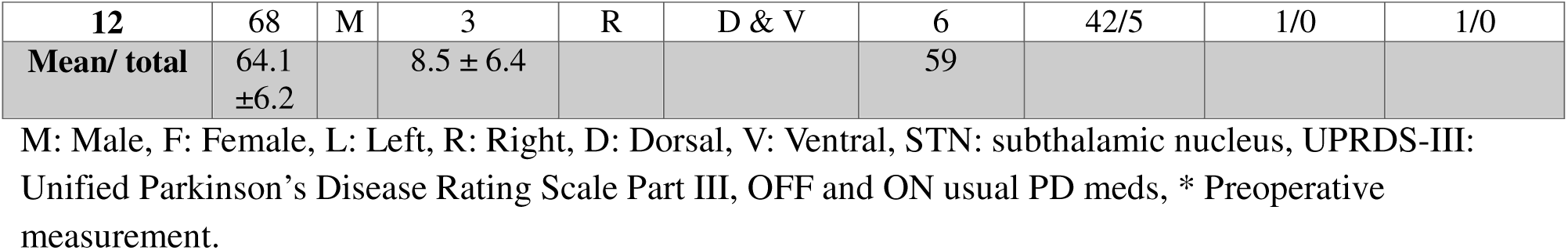
Participant demographics.

| Participant | Age (yrs) | Sex | PD Duration (yrs) | STN Side | STN Subregion Recorded | Number of STN clusters | UPDRS-III Total (Off/On)* | UPDRS-III Speech (Off/On)* | UPDRS-III Facial (Off/On)* |
| --- | --- | --- | --- | --- | --- | --- | --- | --- | --- |
| 1 | 62 | M | 24 | L | D & V | 7 | 37.5/5 | 1/0 | 2/1 |
| 2 | 67 | M | 13 | L | V | 2 | 33/23.5 | 0/1 | 2/1 |
| 3 | 59 | F | 3 | L | D & V | 7 | 31/12 | 1/0 | 1/0 |
| 4 | 64 | M | 10 | R | V | 4 | 43/15 | 1/1 | 1/0 |
| 5 | 73 | M | 4 | R | V | 4 | 38/14 | 0/0 | 1/0 |
| 6 | 69 | M | 7 | R | D | 4 | 43/24 | 1/1 | 1/2 |
| 7 | 56 | M | 2 | L | D & V | 6 | 42/8 | 1/1 | 2/0 |
| 8 | 56 | F | 8 | R | V | 3 | 45/27 | 1/1 | 1/1 |
| 9 | 58 | M | 5 | R | D & D | 3 | 50.5/8 | 2/0 | 2/0 |
| 10 | 74 | F | 16 | R | V | 7 | 38/10 | N/A | N/A |
| 11 | 63 | M | 7 | R | D | 6 | 37/20 | 2/1 | 2/1 |
| 12 | 68 | M | 3 | R | D & V | 6 | 42/5 | 1/0 | 1/0 |
| Mean/ total | 64.1<br>±6.2 |  | 8.5 ± 6.4 |  |  | 59 |  |  |  |

M: Male, F: Female, L: Left, R: Right, D: Dorsal, V: Ventral, STN: subthalamic nucleus, UPRDS-III: Unified Parkinson's Disease Rating Scale Part III, OFF and ON usual PD meds, \* Preoperative measurement.

### Surgical procedure and microelectrode recording

Frame-based stereotactic indirect targeting of STN was utilized and refined by microelectrode recording (MER). Three tracts were made to allow the simultaneous MER from three 2-mm apart center-to-center micro electrodes (∼0.4 to 0.8 MΩ, Sonus, Alpha Omega Inc, Alpharetta, GA, USA) in an anterior / center / posterior arrangement. The dorsal and ventral borders of STN were identified using online analysis of spiking activity conducted with the Neuro Omega recording system (Alpha Omega Inc, Alpharetta, GA, USA). The dorsal border of STN was defined by increased background spiking and bursting activity. Correspondingly, drop-off of spiking activity delineated the ventral border of STN. Once at least one microelectrode was confirmed to be within STN borders, the research protocol was conducted and all microelectrodes were maintained at a fixed depth for the recording. In addition, macrostimulation was performed for clinical reasons after completion of the research protocol to confirm adequate STN localization. Post-operatively the positions of microelectrodes during the research tasks were identified from intraoperative notes based on the depth of the microelectrode relative to the MER-defined STN borders. For this project we divided the STN into dorsal and ventral halves. Of the 36 microelectrode trajectories total, 20 were determined to be in the STN during the research tasks and therefore included in the study.

### Task performance

The behavioral task commenced after clinical MER confirmed intra-STN placement of at least one microelectrode and all intravenous sedatives had been off for at least 30 minutes. The experiment comprised of tasks in two domains: speech and orofacial movement (**Figure 1**). The speech condition included both sentence production and syllable repetition. The orofacial condition included jaw movement (opening and closing) and repetitive tongue protrusion. Together, these orofacial tasks targeted the two principal articulatory motor domains engaged in speech production (Weismer, 2023). For the speech condition, participants were instructed to recite a sentence (“Buy Bobby a Puppy”) followed by a diadochokinetic task (syllable repetition, /tatata/). These tokens were chosen because they are widely used in the assessment of motor speech disorders, including PD, and involve different linguistic and motor complexities (Tjaden, 2008). The sentence was chosen by a speech-language pathologist (SLP; [KT]). The behavioral tasks were self-paced and conducted in two separate blocks in one recording session. The order of task performance was randomized among the participants.

**Figure 1:**
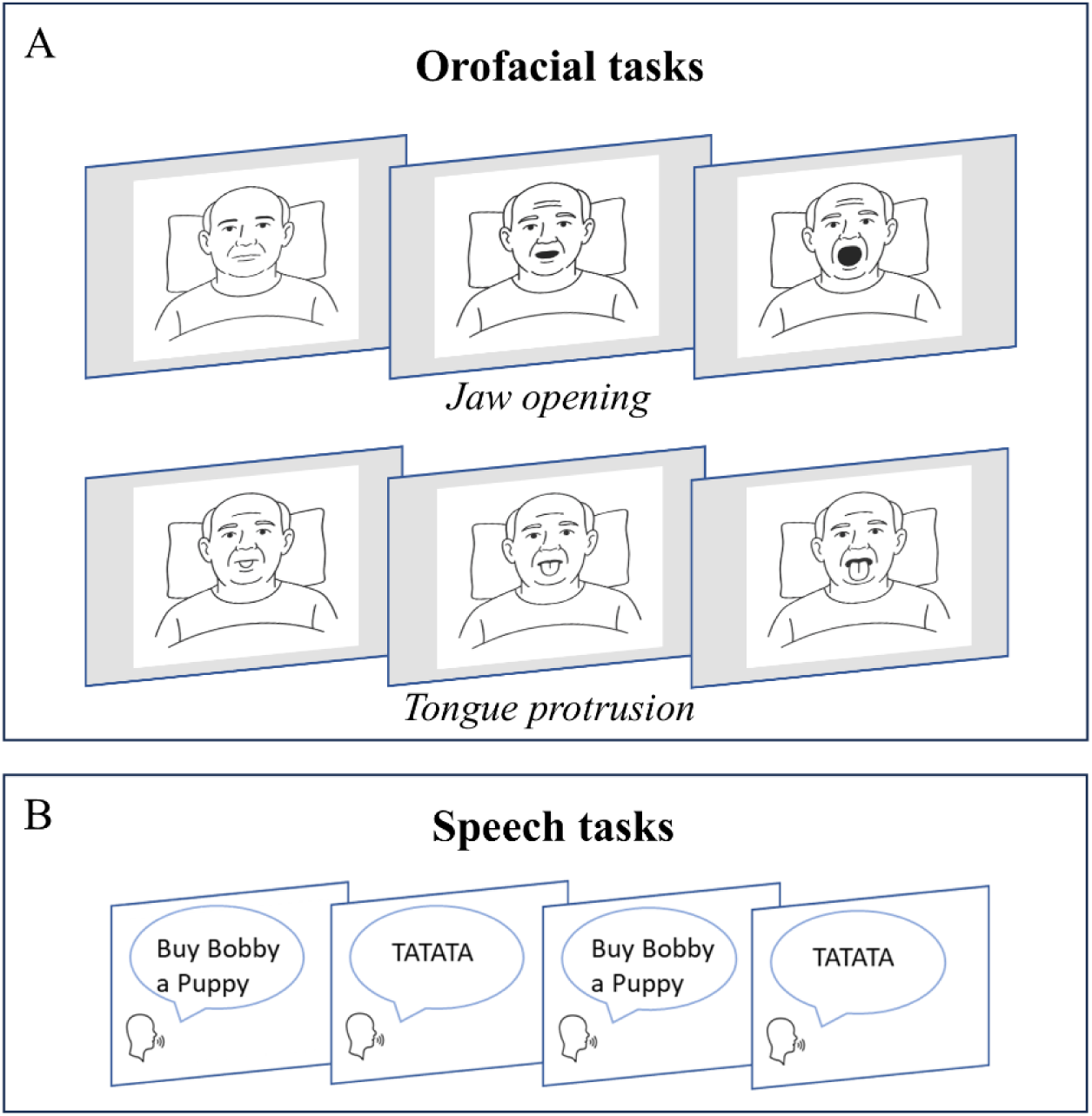
Orofacial (A) and speech (B) tasks performed. A, Participants performed repetitions of self-paced jaw movement (i.e., mouth opening; top row) and tongue protrusion (bottom row) in an interleaved fashion. B, Participants did self-paced, interleaved sentence production (‘Buy Bobby a Puppy’) and syllable repetitions (‘tatata’).

### Data acquisition

Speech was captured via a microphone (E6 omnidirectional microphone, Countryman Associates, Inc., Menlo Park, CA, USA) and sampled at 48 kHz (three participants at 24 kHz). The microphone was fixed in position on the stereotactic frame at an angle of ∼45 degrees, positioned ∼10 cm away from participant’s mouth. The signal was fed into a multi-channel data acquisition system (TDT; Tucker Davis Technologies). The onset and offset of the speech tokens were marked manually using Praat (Boersma & Weenink, 2022). An SLP (AHR) verified the accuracy of the audio timestamps visually.

The orofacial movement tasks were captured via a separate video recorder (24.0 frames / second; Handycam, Sony Corporation, Minato City, Japan). The additional audio recording from this camera was time-aligned with the simultaneously acquired frame-affixed microphone recording to allow a common time scale between the audio and video timeseries with the TDT system. The onsets and offsets of orofacial movements were marked by frame-by-frame visual inspection of the video footage using custom-written MATLAB scripts. The accuracy of the timestamps was verified by a second individual independently.

MER data were passed from the Neuro Omega recording system into the TDT system; therefore all behavioral and electrophysiological data had common timescales for post-hoc analyses. MER were captured at 48 kHz (three participants at 24 kHz) with acquisition filters 0.3 - 9kHz, denoised using the demodulated band transform to filter stationary and non-stationary line noise (C. K. Kovach & Gander, 2016), downsampled to 16 kHz, and high-pass filtered at 200 Hz using a zero-phase finite impulse response (FIR) filter.

### Spike detection and sorting

MER data were sorted offline for single- and multi-unit clusters using an algorithm developed by our lab (C. Kovach et al., 2023). Specifically, filters were estimated for potential candidate unit waveforms through a decomposition of the third-order spectrum, based on the higher-order decomposition method (HOSD) (C. K. Kovach & Howard, 2019). Extracted features were clustered using a mixture-of-Gaussians model in MATLAB, and spike times from these clustered features were used to plot the separate candidate waveforms. Cluster number in the mixture-of-Gaussian’s model was selected as the value that minimized the difference between the observed distribution of distances from cluster centroids and the distribution of distances expected for samples from a multivariate Gaussian distribution. The HOSD approach leverages a recognized characteristic of higher-order spectra, namely the preservation of Fourier-domain phase, by which spike waveform templates and matched filters may be recovered without biases introduced by filtering and thresholding (Shahid et al., 2005). The decompositional approach developed by HOSD improves prior applications of this principle by permitting the recovery of multiple waveforms from a given HOS estimate. The HOSD framework provides spike waveforms and matched filters without the necessity of identifying any candidate spikes beforehand. One advantage of this spike sorting technique is that it can perform substantially better than other spike sorting methods when signal to noise ratio is at intermediate or low levels (C. Kovach et al., 2023).

The identified clusters underwent evaluation to distinguish between single- or multi-unit spiking activity. Single-unit clusters were defined based on the waveforms shape, along with interspike interval distributions that did not generally violate refractory periods and clusters that exhibited distinct, well-separated groupings in PCA space. Any cluster that had more than 2% of their interspike intervals within 2.5 ms and clusters that showed less distinct separation, often indicating the presence of overlapping neural activity or contributions from multiple neurons, were classified as multi-unit.

Neural clusters that were successfully isolated throughout the total length of recording, maintained a firing rate greater than 1 Hz (operationally-defined threshold set to exclude sparsely firing neurons) and did not show regular peaks in the autocorrelogram were selected for further analysis. To visualize and evaluate the spiking rate and timing of STN task-specific neuronal responses, raster plots and spike density functions were generated, created by convolving spike times with a Gaussian kernel having a standard deviation of 20 ms for all events aligned to the onset of each trial **(Figure 2)**. In addition, we examined raw MER data to identify any potential acoustic or voice/microphonic contamination (Berger et al., 2022). STN recordings were first screened by listening for any indications of acoustic contamination. Additionally to screen for acoustic contamination, we applied the method developed by Roussel et al., which quantifies correlations between audio and neural spectrograms across frequencies and time (Roussel et al., 2019). The approach identifies contamination by testing whether correlations are selectively concentrated along the frequency-matched diagonal (i.e., same-frequency coupling) and comparing this pattern against shuffled surrogate data to assess statistical significance. No significant diagonal structure was detected, indicating absence of measurable acoustic contamination in our data.

**Figure 2:**
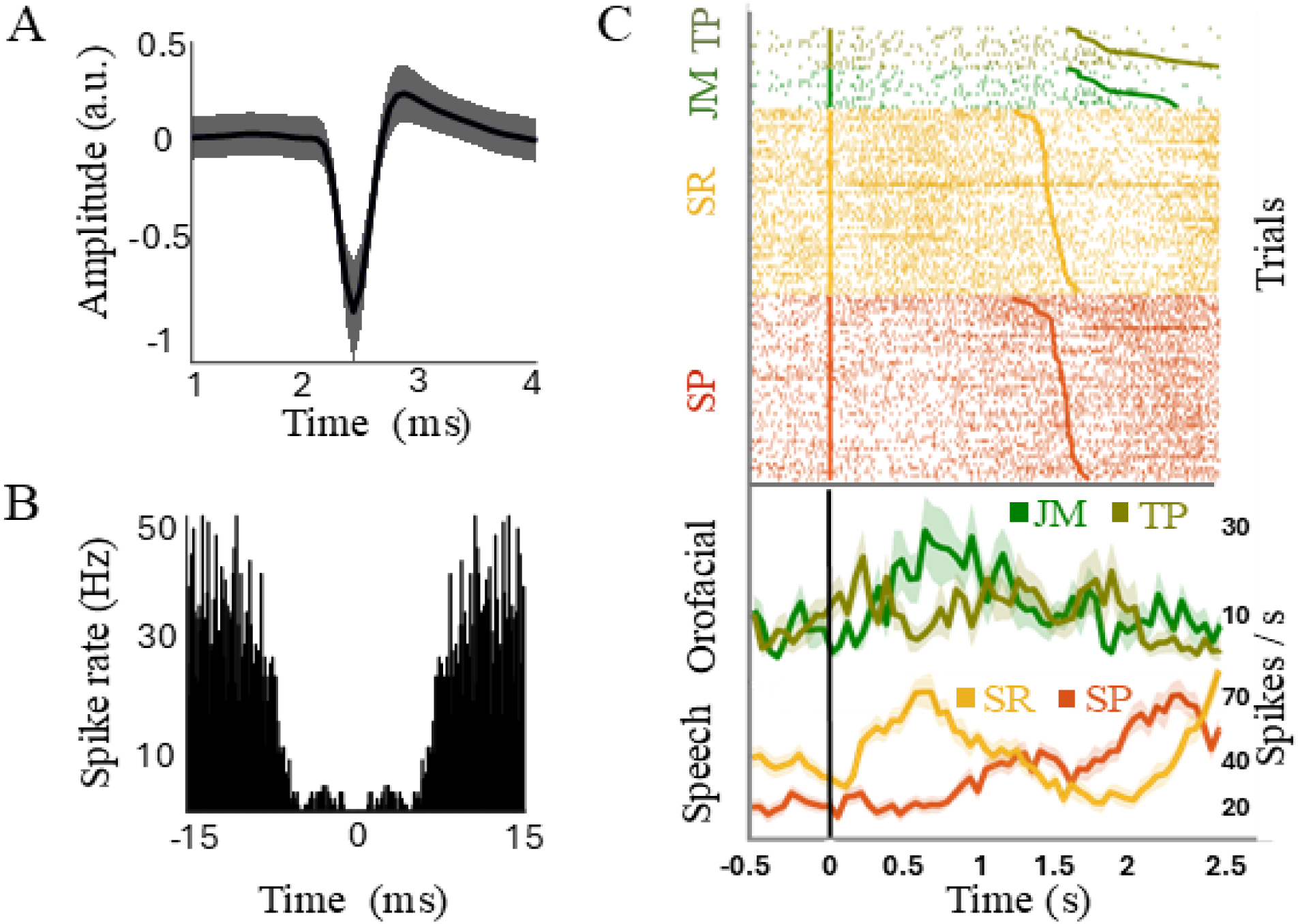
Example STN single-unit response pattern. A, Single-unit extracellular waveform. B, Autocorrelation function indicates the absence of noise peaks and lack of coinciding amplitudes at equal time points suggesting that these waveforms originate from a single unit. C, Raster plots showing spike times of isolated unit for different tasks (upper panels) and average firing rates (lower panel). The vertical line (time 0) denotes task onset, and the smooth line trace represents the offset of each trial, sorted by trial duration. Each orofacial task (green colors) comprised 10 trials, while each speech task (red/orange) included 48 trials. This example neuron exhibited significant modulation during the SR task. JM: jaw movement (kelly green), TP: tongue protrusion (olive green), SP: sentence production (red), SR: syllable repetition (orange).

### Statistical analysis

Previous studies on speech-related STN activity have demonstrated that neural responses vary with different timing aspects of the task. Distinct responses have been observed during the speech planning phase compared to the execution phase (Chrabaszcz et al., 2019; Lipski et al., 2018), as well as at the onset and offset of the speech task (Johari et al., 2023; Watson & Montgomery, 2006). We therefore developed a generalized linear model (GLM) using a Poisson rate function, to specify the probability of average spiking activity over binned time intervals as a function of various covariates. These covariates represented different temporal phases of the task: 1) pre-task (from 300 ms before to task onset) versus task execution, consistency across trials (from initial trials to final trials) for 2) total trial interval (including pre- and post-task execution time) and 3) task execution only (onset to offset), and 4) firing changes over time within the trials. This model fitting procedure follows a similar approach to one used by others (Kramer & Eden, 2016; Truccolo et al., 2005).

Spiking activity was assumed to be a discrete time point process and follow a Poisson distribution. The GLM was formulated as:

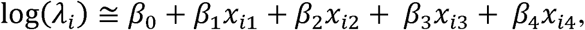

where *λ_i_* is the spike counts in time bin *i*, *β*_0_ to *β*_4_ are regression coefficients, and *x_i_*_1_ to *x_i_*_4_ represent the 4 covariates designed to account for time-dependent aspects of the tasks (**Figure 3**). The estimated coefficients quantified the direction of the relationship between the covariates and changes in spike counts.

**Figure 3.**
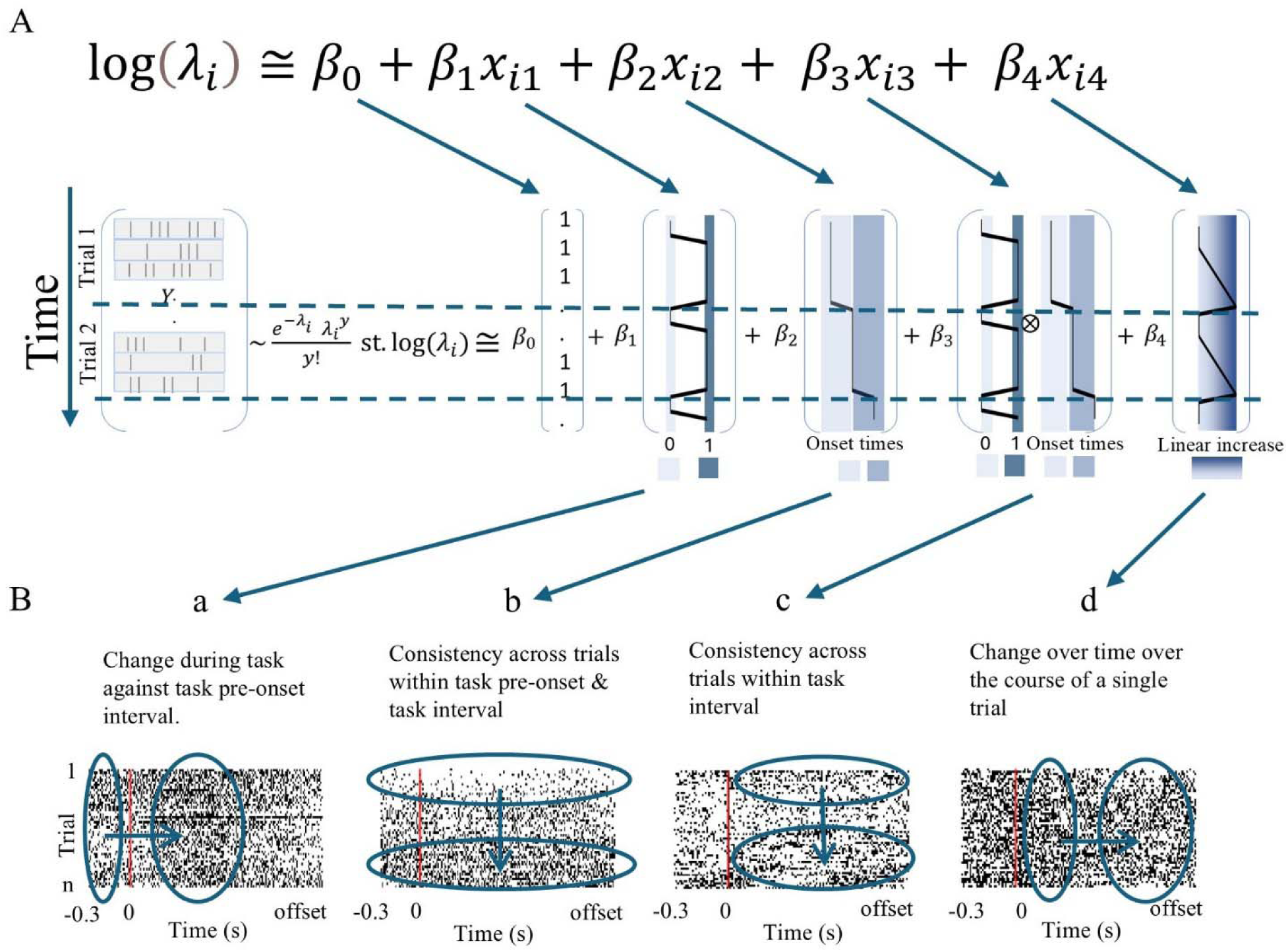
A, GLM design. Response variable (Y) is spike counts in sequential 10 ms bins. Y is assumed to follow a Poisson distribution with its mean estimated as a linear function of 4 variables where the significance of the fitted corresponding coefficient (β) indicates the responsiveness of the neural cluster to the task. In the first covariate, 0 (light blue) and 1 (dark blue) represent pre-onset intervals and task intervals respectively. Second covariate linearly increases as trials advance from first trial (light blue) to subsequent trials (dark blue). The third covariate represents the interaction between the first and second covariates, capturing trial progression exclusively during task execution. Fourth covariate captures any linear trend over time within the trial window. Light blue represents the onset and darkest blue indicates the offset. B, Example clusters of statistically significant coefficients. Subpanel a illustrates changes in the spike rate intensity from the pre-onset interval to the task interval. Subpanel b represents the changes in the spike rate intensity over trials. Subpanel c demonstrates changes in spike activity over trials during the task execution period only. Subpanel d depicts changes in spike rate intensity with linear time increase within a trial.

Covariate 1, *x_i_*_1_, accounted for the task relative to pre-task interval (i.e., trial-by-trial total task duration vs. total pre-task interval). This means *x_i_*_1_ takes the value of 1 during the task execution window and 0 during the 300 ms long pre-task window (**Figure 3-Ba**). Covariate 2, *x_i_*_2_, characterized the trial duration across baseline and task interval to account for any task-independent linear trends over the course of the trials (i.e., early vs later trials). This means *x_i_*_2_ takes the same value throughout pre-task and task window for each trial (set as the task onset time) and linearly increases as the trials progress. To remove the correlation with the intercept, *x_i_*_2_ was mean-centered. It is important to note that this covariate was not utilized to demonstrate task-related neural modulation, rather it was designed to regress out its effect on the other covariates. This covariate therefore controlled for confounding factors such as global effects or non-task-related neural modulations (**Figure 3-Bb**). Covariate 3, *x_i_*_3_, captured the effect of trial progression during task execution interval (i.e., total trial duration from onset to offset) over time, as the interaction between Covariates 1 and 2 (**Figure 3-Bc**). Covariate 4, *x_i_*_4_, accounted for the linear progression of time within the trial interval from the trial onset to the trial offset. This covariate described each trial by the same linear pattern. Hence, it captured any cumulative response of the neurons throughout the duration of the task **(Figure 3-Bd**).

To capture rapid changes in neural activity, trials were segmented into 10 ms bins. The GLM was fit separately for each cluster and each task using the function glmfit in MATLAB, with the spike count as the dependent variable and condition as the independent variable. Four covariates were included to represent different temporal aspects of task performance. To evaluate model performance, we compared a null model containing only the intercept to the full four-covariate model using likelihood ratio p-value (pllr). We used Akaike Information Criterion (AIC) to assess whether inclusion of the additional covariates significantly improved fit beyond covariate 2 which captures global effects or non-task-related neural modulations.

Task specific cluster modulation was assessed by testing the significance of each covariate in the GLM. Clusters with at least one significant covariate (Covariates 1, 3, 4) were classified according to the sign of the corresponding regression coefficient and according to their modulation profile as: increasing, where the covariate exhibited a positive correlation with spike counts (indicating that higher values of the covariate correspond to increased firing rates); decreasing, where the covariate displayed a negative correlation with spike counts (indicating that higher values of the covariate correspond to reduced firing rates); or mixed, which was assigned when multiple covariates were significant but their coefficients suggested opposing effects (i.e., some covariates positively and others negatively influencing firing rate).

For correlation analyses evaluating relationships between spontaneous STN firing rates and clinical measures, a baseline firing rate was calculated for each unit. The spontaneous baseline was defined as the average spike rate during a period from −500 to −200 ms prior to behavioral event onset. To minimize contamination from preceding trials, only trials with inter-trial intervals greater than 700 ms were included in this calculation.

To compare means, we performed Wilcoxon rank and Kruskal-Wallis tests. We used Spearman’s correlations to measure the degree of dependence between numerical and ordinal variables and Pearson’s correlations to measure the relationship between two numerical variables. To assess significance of the difference between the frequency distribution of the categorical variables, we used a chi-square test of independence. All analyses were conducted in MATLAB 2022a (R2022a, Mathworks Inc), and a significance threshold for all analyses was set at α=0.05. All p-values were adjusted using false discovery rate for multiple comparisons where applicable.

## Results

The 12 participants performed on average 39 ± 11 (mean ± std) repetitions of sentence production, 38 ± 11 trials of syllable production, 8 ± 3 trials of repetitive jaw movement, and 7 ± 2 trials of repetitive tongue protrusion tasks in two separate blocks. Following curation of neural recordings to retain well-isolated units we identified a total of 59 single- and multi-unit STN clusters. Between two and six clusters were identified per microelectrode. The proportion of detected clusters per electrode was not correlated with PD duration (*p*=0.97, *r*=0.01, Spearman correlation) or severity (i.e., higher UPDRS-III Off score; *p*=0.23, *r*=-0.37, Spearman correlation). Clusters that did not show consistent firing rate above 1 Hz throughout the recording (2 clusters) and clusters that were only present during one block (6 clusters) were excluded from further analysis. Ultimately, 51 clusters comprising 23 single-units and 28 multi-unit neural clusters were present throughout the entire recording duration and included in the final analysis. The mean spontaneous baseline firing rates were 14 ± 7 spikes/s and 37 ± 14 spikes/s for single- and multi-unit clusters respectively. These values fall within the range of the baseline firing activity reported in prior studies involving single- and multi-unit clusters isolated from the human STN (Remple et al., 2011; Watson & Montgomery, 2006). The mean baseline firing rates were slightly higher in clusters that showed response to speech tasks (33 ± 22 spikes/s) compared to those that were not significantly modulated during the speech task (25 ± 14 spikes/s). However, this difference did not reach statistical significance.

Across all model fits, 67% demonstrated a significant neural response to the tasks, as indicated by the likelihood ratio test comparing the full four-covariate model to the intercept-only null model (p < 0.05). The full model outperformed the reduced model (intercept + covariate 2 only) in 59% of cases (AIC comparison). A cluster was classified as responsive / modulated by a task if at least one covariate (Covariates 1, 3,4) achieved statistical significance in the GLM. Overall, 31/51 clusters (61%) showed a response to either of the tasks, with 11 clusters (22%) responding to the speech tasks, 6 clusters (12%) to the orofacial tasks, and 14 clusters (27%) to both tasks (**Figure 4A**). Varied response types were found, with some clusters showing increased, decreased, or mixed directions of firing rate changes (**Figure 4A**, left bars). In speech-exclusive responses, increased and decreased firing rates occurred in equal proportions (50% each), whereas orofacial-exclusive responses showed a higher proportion of decrease in firing rates (67%). Clusters responsive to both task types more often showed increased firing rates (47%). For specific covariate response patterns, most task-related modulations were attributed to significant responses in a single covariate. However, eight task-related modulations resulted from significant responses in two covariates, while one task-related modulation was from significant responses in three covariates.

**Figure 4.**
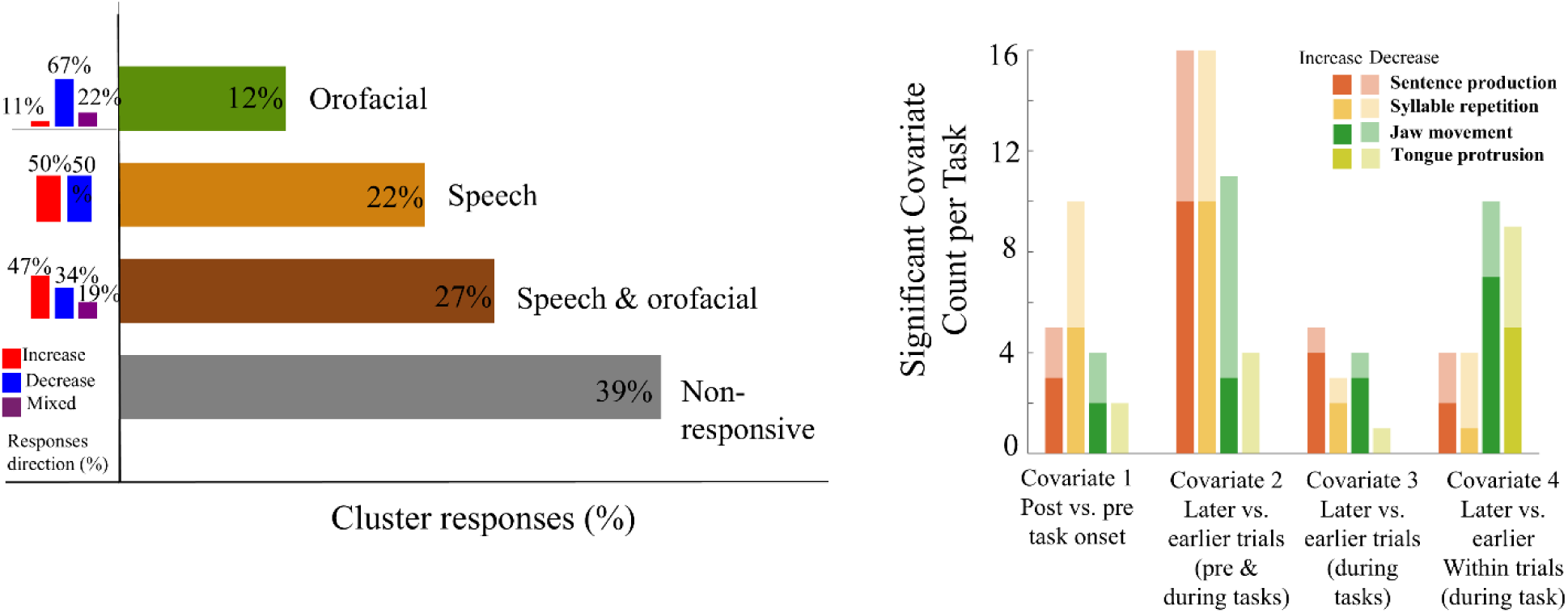
A, Frequency of task responsive and non-responsive clusters. The bar plots to the left show the direction of responses. B, The frequency and direction of significant covariates. The total number of significant clusters by covariate is indicated by the column height, and the stacked colors show the proportion of increased (higher color intensity) and decreased (lower color intensity) firing rate changes are shown for each task and each covariate.

**Figure 4B** shows significant clusters by covariate, as well as the direction of firing rate change (increased or decreased). Changes in firing rates during the task versus pre-task onset (Covariate 1) were more frequently observed in neurons during speech tasks compared to orofacial tasks (15 vs. 6 clusters). Increased firing in Covariate 1 means higher firing rate in the post-onset compared to pre-onset periods. Changes in firing rate across the later compared to earlier trials (Covariate 2) were found in 32 speech-related clusters and 15 orofacial clusters, with a greater proportion showing increased firing over time. During the task period, the frequency of firing rate changes from later trials to earlier trials (Covariate 3) was more frequent in speech than orofacial tasks (8 vs. 5 clusters). In contrast, firing rate changes from later to earlier in each trial during the task interval (Covariate 4) were more prevalent in orofacial tasks than in speech tasks (19 vs. 8) and were more often reflected in increased rather than decreased firing rates.

We found that 12% of clusters (6/51) responded only to the sentence production task, 10% (5/51) only to syllable repetition, 2% (1/51) only to jaw movement, and 4% (2/51) only to tongue protrusion (**Table 2**). Co-occurrence of significant responses among the task subtypes was also observed. The co-occurrence of syllable repetition and sentence production was the least frequent (2%), whereas the co-occurrence of syllable repetition and jaw movement was the most frequent (15%). The co-occurrence of jaw movement and tongue protrusion was the second most frequent (12%).

**Table 2.** Response occurrences.

|  | Sentence production | Syllable repetition | Jaw movement | Tongue protrusion |
| --- | --- | --- | --- | --- |
| Sentence production | 12% * | - | - | - |
| Syllable repetition | 2% | 10% * | - | - |
| **Jaw movement** | 8% | 15% | 2%* | - |
| **Tongue protrusion** | 6% | 8% | 12% | 4%* |

Population-level significance was tested with binomial tests comparing the observed fraction of responsive neurons to the expected chance level, derived from the neuron-level FDR-corrected threshold (Benjamini–Hochberg, *q* < 0.05). The proportion of task modulated clusters was significantly higher than chance for sentence production (Binomial test, chance level = 0.01, *p* < 0.001) and syllable repetition tasks exclusively (Binomial test, chance level = 0.01, *p* < 0.001) but not for jaw movement (Binomial test, chance level = 0.01, *p* = 0.40) or tongue protrusion exclusively (*p* = 0.09). While the proportion of clusters modulated by both sentence production and syllable repetition tasks was not significant (Binomial test, chance level = 0.01, *p* = 0.40), the proportion of task modulated clusters to paired jaw movement and tongue protrusion was significant (Binomial test, chance level = 0.01, *p* < 0.001). Neural cluster modulations to the combination of speech and orofacial tasks were significant, with 27% (14/51) of clusters showing responsiveness (Binomial test, chance level = 0.01, *p* < 0.001).

### Correlation between the neural activity and MER topography

Electrode recording locations within STN were distributed equally between the dorsal and ventral halves (9 vs. 10 locations). There was no correlation between recording depth (expressed as the percentage of MER depth from the dorsal border of the STN) and either the proportion of task-specific cluster modulation (Kruskal–Wallis test, *p* = 0.35) or the proportion of unit modulation type direction (i.e., increase or decrease in firing rate; Kruskal–Wallis test, *p* = 0.58). A higher proportion of non-task-modulated neural clusters was observed at the dorsal and ventral borders of the STN (55%) compared to modulated clusters (45%). In contrast, within the middle portion of the STN, task-modulated clusters were more prevalent (71% vs. 29%). However, this difference did not reach statistical significance [χ² (1, N=51) = 3.43, p = 0.06] and recording locations were heavily skewed to the middle of STN **(Figure 5).**

**Figure 5.**
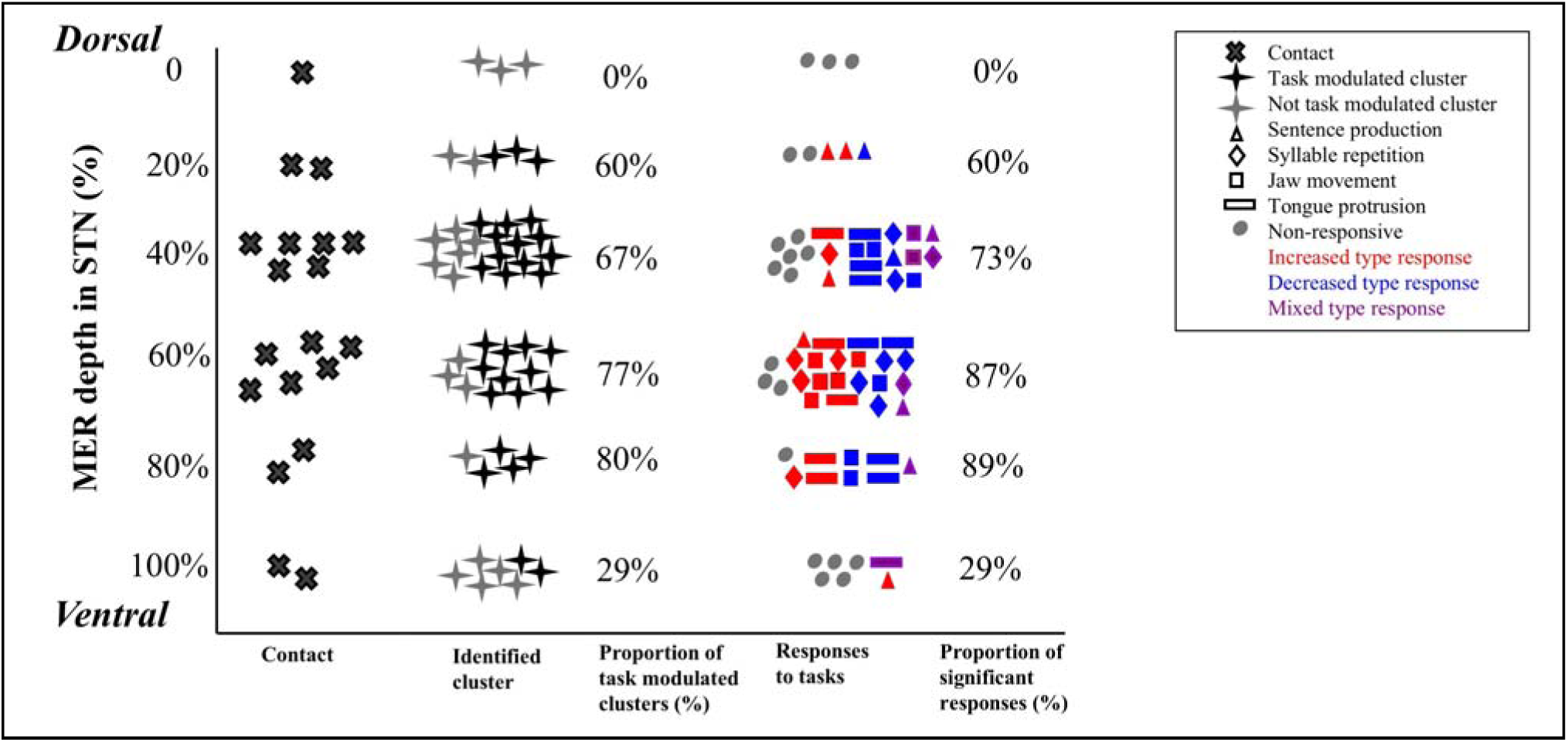
STN recording location distribution (first column) and identified clusters (second column). Recording locations were predominantly located in the middle portion of STN. The percentage of task modulated clusters to total identified clusters is shown in the center column. Not modulated clusters were seen to lie more commonly closer to dorsal and ventral STN borders. The response pattern by task is shown in the fourth column. Task type is denoted by different shapes. Response type is indicated by color: increased firing rates are red, decreased are blue, and mixed firing changes are purple. The proportion of significant responses is depicted in the right column.

There were more electrode recording locations in the right STN than in the left (12 vs. 7), and a greater number of neural clusters were identified in the right STN (34 vs. 17). Overall, the proportion of task modulated neurons was higher on the left side (76%) compared to right (53%). However, a Chi-square test revealed no significant correlation between recording hemisphere (right vs. left) and task-specific cluster modulation [χ² (3, N=51) = 4.05, p = 0.25]. The proportion of modulated clusters responding to speech, orofacial, and both tasks was 15%, 15%, and 23% in the right STN, respectively, compared to 35%, 12%, and 29% in the left STN.

Participants who had higher levodopa responsiveness, defined as UPDRS-III total improvement score, calculated as (UPDRS-III total score OFF PD medications - UPDRS-III total score ON PD medications) / UPDRS-III total OFF score, had a higher proportion of non-responsive neural clusters (p<0.001, Wilcoxon signed rank test). There was no correlation between UPDRS-III speech subscale OFF score or UPDRS-III orofacial subscale OFF score and proportion of speech responsive or orofacial responsive neural clusters, respectively (rho= -0.03, *p*=0.92 and rho=0.06, *p*=0.85, Spearman correlation). PD duration was not correlated with the proportion of responsive neural clusters (rho= -0.45, *p*=0.15, Spearman correlation). Additionally, the spontaneous firing rate showed no significant correlation with PD duration (rho=-0.24, *p*=0.09, Pearson correlation) or disease severity, as measured by the total OFF score of the UPDRS-III (rho= -0.25, *p*=0.08, Pearson correlation).

## Discussion

Speech is a sensorimotor process in which cognitive and linguistic intentions are transformed into precise articulatory movements under strict temporal constraints, engaging multiple muscle groups to generate the diverse patterns of spoken language (Hickok et al., 2003; Simmonds et al., 2014). Previous research has shown robust STN neural modulation during speech tasks, with 38–79% of recorded neurons exhibiting speech-related modulation across studies (Johari et al., 2023; Lipski et al., 2018, 2024; Tankus & Fried, 2019a; Watson & Montgomery, 2006). These findings highlight the STN as a subcortical node in speech-related processes. Analyses of single- and multi-unit neural activity as well as local field potentials suggest that such modulation may reflect contributions from motor-related mechanisms, e.g. planning (Lipski et al., 2018), execution (Lipski et al., 2018), respiratory-phonatory control (Chrabaszcz et al., 2019), linguistic processes (e.g., syntactic structure and phonetic content (Lipski et al., 2024)), sensory-motor integration (Vissani et al., 2025), and sensory-timing functions (Lipski et al., 2018).

Here, we aimed to investigate the speech-specificity of neural modulation in STN. By analyzing direct STN recordings across different temporal windows of speech and non-speech orofacial tasks, we sought to better understand the role of STN in speech production. Our results revealed a higher proportion of STN neural clusters were modulated (61%) than not (39%) by both speech and orofacial tasks. The largest proportion of clusters were modulated by both speech production and orofacial movement (27%), suggesting a shared STN substrate for these tasks. However, STN modulation to speech tasks alone compared to orofacial tasks alone was common (22% vs. 12%). The greater modulation during speech production suggests that STN activity during speech is driven by mechanisms beyond simple orofacial movements necessary for speech, likely reflecting additional cognitive and linguistic processes in STN.

Covariate 1 compared spike counts between a pre-task interval and task execution, thereby studying neural modulation across both preparatory and performance phases. Clusters that change activity prior to task onset may reflect the STN’s role in either motor alone or speech motor planning, whereas those that show spike count changes during execution align more closely with active motor output. We observed a greater number of clusters showing significant modulation during the task compared to the pre-task interval for speech compared to orofacial tasks (15 vs. 6 clusters), suggesting speech-specificity of STN modulation. Increases in spike counts during the task compared to the pre-task interval were slightly more prevalent (8/15 clusters) compared to decreased firing rates. Conversely, decreased spike counts between these time periods were more common for the orofacial tasks (4/6 clusters). Other reports have shown STN modulation from –180 ms to +50 ms relative to speech onset and related the changes in firing rate during a speech task to motor execution processes (Lipski et al., 2018). Supporting this view, Chrabaszcz et al. (Chrabaszcz et al., 2019) showed that STN high-gamma activity (as a proxy for firing rate) increased before and persisted throughout articulation, linking the STN to both planning and execution phases of speech production. Together, these results suggest that Covariate 1 effectively captured the dual role of STN neurons in both planning and executing speech and orofacial tasks, with a particularly prominent contribution during speech.

Repetitive motor control arises from dynamic interactions between cortical planning and feedback systems and subcortical circuits that fine-tune performance across successive repetitions (Rogalsky et al., 2015; Seidler et al., 2004). Covariate 3 also captured processes related to task repetition in STN by assessing trial-progression effects but only during the task execution period. In our dataset, a comparable number of clusters showed significant spike count changes of later versus earlier trials for both speech and orofacial tasks (8 vs. 5 clusters) and ratios of increased versus decreased spike counts (6/8 vs 3/5 respectively). These findings suggest that STN adaptation is a general mechanism supporting repetitive motor tasks rather than a speech-specific process. Most speech related studies in STN have focused on firing-rate changes during or immediately before task execution, but little is known about how STN activity evolves across repeated trials so follow-up studies are needed.

We examined within-trial spike counts dynamics (Covariate 4) to track changes in firing rates from task onset to the completion of individual trials. Previous studies have shown that STN neurons exhibit diverse modulation patterns time-locked to both the onset and offset of speech tasks (Johari et al., 2023; Lipski et al., 2024). As highlighted by Lipski et al. (Lipski et al., 2018), such variability may reflect ongoing monitoring of task performance. In our dataset, we observed a greater number of clusters modulated later compared to earlier in the trial for orofacial compared to speech tasks (19 vs. 8 clusters). Over the course of the trials, the orofacial spike count changes were more often increased (12/19 clusters) than found during the speech tasks (3/8 clusters). This temporal pattern difference suggests speech-specific differential monitoring mechanisms in STN during task execution. Speech production requires coordination of linguistic, cognitive, and sensorimotor integration processes, therefore larger datasets are required to understand and parse STN dynamics between these subprocesses.

The strongest paired co-occurrence of significant STN modulations (15%) was identified between clusters modulated by syllable repetition and jaw movement, suggesting a shared motor pathway due to their comparable articulatory requirements. Conversely, the least frequent co-occurrence was between sentence production and syllable repetition. However, both sentence production (12%) and syllable repetition (10%) demonstrated prominent modulation independently, further supporting a speech-specific role of STN.

Our data revealed varying directions of STN neural modulation, via increased, decreased, or mixed changes in task-specific firing rates. As a critical component of a cortico-basal ganglia-thalamo-cortical circuit, STN integrates both excitatory and inhibitory inputs and is known to be engaged in both motor and cognitive functions (Gradinaru et al., 2009; Temel et al., 2005). Consequently, it is not surprising that we observed a diverse pattern in the direction of firing rate changes in the STN with our tasks. In addition, our findings align with the results of previous studies indicating mixed STN response to speech and motor tasks (Johari et al., 2023; Lipski et al., 2018).

Historically, the human STN has been described as comprising three functional subdivisions: a dorsal one-third primarily associated with motor functions, a middle one-third linked to cognitive and associative processes, and a ventral one-third related to limbic and cognitive functions (Mallet et al., 2007; Temel et al., 2005). While we did have some data from each of these thirds, our data was heavily skewed to the mid-portion of STN and therefore underpowered to report subregional differences with STN. The mid-portion of the STN contained a high proportion of task-modulated neurons, consistent with reports that projections from primary and secondary motor cortical areas converge on central STN regions (Jeon et al., 2022). In our limited number of clusters in ventral STN, we did observe modulation by our tasks. Together, these preliminary findings suggest that while broad functional gradients may exist across the STN, its neural activity during speech and orofacial tasks is not strictly confined to classical motor–associative–limbic subdivisions (Prasad & Wallén-Mackenzie, 2024).

We did not find a correlation between the duration or severity of PD and spontaneous STN baseline firing rates or proportion of modulated clusters. Some studies have reported an increased baseline firing rate in the human STN with worsening disease severity (Johari et al., 2023). Studies conducted on primates have also demonstrated a significant increase in STN firing rate with dopamine depletion in the STN (Bergman et al., 1990). Our data did show a correlation between greater levodopa responsiveness (i.e., ON/OFF improvement) and non-modulated STN clusters. Additional study is needed for further insight into the relation between PD disease features and the firing rate of STN neurons.

Our study was not designed or powered to investigate hemispheric differences in STN during speech production. More clusters were captured in the right than left STN and the slightly higher rate of task-modulated responses in the left was not statistically significant, possibly due to the overall number of clusters captured. Our study, along with others such as Tankus et al. (Tankus & Fried, 2019b) and Johari et al. (Johari et al., 2023), has demonstrated STN neural modulation to speech tasks in both the right and left STN. Thus, it can be anticipated that bilateral STN play a role in speech production. Hyder et al. similarly showed involvement of both STN in speech production, albeit in different aspects of speech production (Hyder et al., 2021). There are prior studies suggesting hemispheric laterality in the STN during speech production. Studies by Wang et al (Wang et al., 2006) and Santes et al (Santens et al., 2003) showed a stronger impact of the left STN stimulation on speech outcomes. Further investigations are warranted to elucidate the interhemispheric disparities in STN involvement in speech production.

## Caveats and future directions

As in other human intraoperative STN studies, our analyses were constrained by the number of neuronal clusters (n = 51) that could be reliably held across the duration of the tasks. Additional studies are therefore needed to augment our findings. By necessity we utilized simple speech and movement tasks and these may not fully engage or reflect the role of STN in more complex or longer lasting speech and movement tasks. In particular, more cognitively demanding speech tasks may demonstrate different modulation of STN for planning, initiation, and control of ongoing tasks. Another limitation of this study is the unbalanced number of trials across tasks likely affected the extent of observed modulation. Finally, our findings relate to PD subjects and may not reflect STN modulation in other patient populations.

## Conclusion

Collectively, these findings show the involvement of STN in both speech-specific and overlapping speech and oromotor control functions. This finding extends previous studies showing a role of STN in speech production. While this supports STN’s role in integrating motor and cognitive components of speech production, further research is needed to determine its contribution to more complex language domains and to clarify its interactions with cortical and subcortical speech and language related networks.

## Funding resources

This project is supported by NIH grant R01DC017718-01A1.

## Competing interest

Authors declare no competing interest.

## Acknowledgements

We sincerely thank the patients and their families for generously contributing their time and effort to this study. We also gratefully acknowledge Haiming Chen for assistance with data collection.

## CRediT

**Zahra Jourahmad:** Conceptualization, methodology, software, validation, formal analysis, data curation, writing original draft, visualization, project administration, writing-review and editing.

**Christopher K. Kovach:** Software, methodology, validation, writing-review and editing.

**Andrea H. Rohl:** Investigation, validation, writing-review and editing.

**Joel I. Berger:** Writing-review and editing.

**Kris Tjaden:** Methodology, project administration, funding acquisition, writing-review and editing.

**Jeremy D.W. Greenlee:** Conceptualization, methodology, supervision, project administration, funding acquisition, writing-review and editing.

## Notes

### Competing Interest Statement

The authors have declared no competing interest.

## References

Aldridge, D., Theodoros, D., Angwin, A., & Vogel, A. P. (2016). Speech outcomes in Parkinson’s disease after subthalamic nucleus deep brain stimulation: A systematic review. Parkinsonism & Related Disorders, 33, 3–11. 10.1016/j.parkreldis.2016.09.022

Atkinson-Clement, C., Maillet, A., LeBars, D., Lavenne, F., Redouté, J., Krainik, A., Pollak, P., Thobois, S., & Pinto, S. (2017). Subthalamic nucleus stimulation effects on single and combined task performance in Parkinson’s disease patients: A PET study. Brain Imaging and Behavior, 11(4), 1139–1153. 10.1007/s11682-016-9588-4

Benabid, A. L., Chabardes, S., Mitrofanis, J., & Pollak, P. (2009). Deep brain stimulation of the subthalamic nucleus for the treatment of Parkinson’s disease. The Lancet Neurology, 8(1), Article 1. 10.1016/S1474-4422(08)70291-6

Berger, J. I., Johari, K., Kovach, C. K., & Greenlee, J. D. (2022). Speech artifact is also present in spike data. NeuroImage, 263, 119642. 10.1016/j.neuroimage.2022.119642

Bergman, H., Wichmann, T., & DeLong, M. R. (1990). Reversal of experimental parkinsonism by lesions of the subthalamic nucleus. *Science (New York*, N.Y.), 249(4975), Article 4975. 10.1126/science.2402638

Boersma, P., & Weenink, D. (2022). Praat (Version 6.2.13) [Computer software]. https://www.fon.hum.uva.nl/praat/

Chrabaszcz, A., Neumann, W.-J., Stretcu, O., Lipski, W. J., Bush, A., Dastolfo-Hromack, C. A., Wang, D., Crammond, D. J., Shaiman, S., Dickey, M. W., Holt, L. L., Turner, R. S., Fiez, J. A., & Richardson, R. M. (2019). Subthalamic Nucleus and Sensorimotor Cortex Activity During Speech Production. Journal of Neuroscience, 39(14), Article 14. 10.1523/JNEUROSCI.2842-18.2019

Deuschl, G., Schade-Brittinger, C., Krack, P., Volkmann, J., Schäfer, H., Bötzel, K., Daniels, C., Deutschländer, A., Dillmann, U., Eisner, W., Gruber, D., Hamel, W., Herzog, J., Hilker, R., Klebe, S., Kloß, M., Koy, J., Krause, M., Kupsch, A., … Voges, J. (2006). A Randomized Trial of Deep-Brain Stimulation for Parkinson’s Disease. New England Journal of Medicine, 355(9), Article 9. 10.1056/NEJMoa060281

Friederici, A. D. (2011). The Brain Basis of Language Processing: From Structure to Function. Physiological Reviews, 91(4), Article 4. 10.1152/physrev.00006.2011

Gradinaru, V., Mogri, M., Thompson, K. R., Henderson, J. M., & Deisseroth, K. (2009). Optical Deconstruction of Parkinsonian Neural Circuitry. Science, 324(5925), 354–359. 10.1126/science.1167093

Guenther, F. H. (2016). Neural Control of Speech. The MIT Press.

Guenther, F. H., & Vladusich, T. (2012). A Neural Theory of Speech Acquisition and Production. Journal of Neurolinguistics, 25(5), Article 5. 10.1016/j.jneuroling.2009.08.006

Hickok, G., Buchsbaum, B., Humphries, C., & Muftuler, T. (2003). Auditory–Motor Interaction Revealed by fMRI: Speech, Music, and Working Memory in Area Spt. Journal of Cognitive Neuroscience, 15(5), 673–682. 10.1162/jocn.2003.15.5.673

Hyder, R., Højlund, A., Jensen, M., Johnsen, E. L., Østergaard, K., & Shtyrov, Y. (2021). STN-DBS affects language processing differentially in Parkinson’s disease: Multiple-case MEG study. Acta Neurologica Scandinavica, 144(2), Article 2. 10.1111/ane.13423

Janssen, N., & Mendieta, C. C. R. (2020). The Dynamics of Speech Motor Control Revealed with Time-Resolved fMRI. Cerebral Cortex, 30(1), 241–255. 10.1093/cercor/bhz084

Jeon, H., Lee, H., Kwon, D.-H., Kim, J., Tanaka-Yamamoto, K., Yook, J. S., Feng, L., Park, H. R., Lim, Y. H., Cho, Z.-H., Paek, S. H., & Kim, J. (2022). Topographic connectivity and cellular profiling reveal detailed input pathways and functionally distinct cell types in the subthalamic nucleus. Cell Reports, 38(9), Article 9. 10.1016/j.celrep.2022.110439

Johari, K., Kelley, R. M., Tjaden, K., Patterson, C. G., Rohl, A. H., Berger, J. I., Corcos, D. M., & Greenlee, J. D. W. (2023). Human subthalamic nucleus neurons differentially encode speech and limb movement. Frontiers in Human Neuroscience, 17. https://www.frontiersin.org/articles/10.3389/fnhum.2023.962909

Jorge, A., Lipski, W. J., Wang, D., Crammond, D. J., Turner, R. S., & Richardson, R. M. (2021). *Hyperdirect connectivity of opercular speech network to the subthalamic nucleus* (p. 2021.07.02.450909). bioRxiv. 10.1101/2021.07.02.450909

Kovach, C., Carvalho Moreira, J., Berger, J., III, M., Gwilliams, L., Fallah, A., Comstock, L., & Mendes, E. (2023). Higher-Order Spectral Decomposition Applied to Spike Sorting. 10.13140/RG.2.2.28352.40967

Kovach, C. K., & Gander, P. E. (2016). The demodulated band transform. Journal of Neuroscience Methods, 261, 135–154. 10.1016/j.jneumeth.2015.12.004

Kovach, C. K., & Howard, M. A. (2019). Decomposition of higher-order spectra for blind multiple-input deconvolution, pattern identification and separation. Signal Processing, 165, 357–379. 10.1016/j.sigpro.2019.07.007

Kramer, M. A., & Eden, U. (2016). Case Studies in Neural Data Analysis A Guide for the Practicing Neuroscientist. MIT Press.

Lipski, W. J., Alhourani, A., Pirnia, T., Jones, P. W., Dastolfo-Hromack, C., Helou, L. B., Crammond, D. J., Shaiman, S., Dickey, M. W., Holt, L. L., Turner, R. S., Fiez, J. A., & Richardson, R. M. (2018). Subthalamic Nucleus Neurons Differentially Encode Early and Late Aspects of Speech Production. The Journal of Neuroscience, 38(24), 5620–5631. 10.1523/JNEUROSCI.3480-17.2018 https://doi.org/10.1101/2023.12.11.569290

Lipski, W. J., Bush, A., Chrabaszcz, A., Crammond, D. J., Fiez, J. A., Turner, R. S., & Richardson, R. M. (2024). Subthalamic nucleus neurons encode syllable sequence and phonetic characteristics during speech. Journal of Neurophysiology, 132(5), 1382–1394. 10.1152/jn.00471.2023

Mallet, L., Schüpbach, M., N’Diaye, K., Remy, P., Bardinet, E., Czernecki, V., Welter, M.-L., Pelissolo, A., Ruberg, M., Agid, Y., & Yelnik, J. (2007). Stimulation of subterritories of the subthalamic nucleus reveals its role in the integration of the emotional and motor aspects of behavior. Proceedings of the National Academy of Sciences, 104(25), 10661–10666. 10.1073/pnas.0610849104

Manes, J. L., Bullock, L., Meier, A. M., Turner, R. S., Richardson, R. M., & Guenther, F. H. (2024). A neurocomputational view of the effects of Parkinson’s disease on speech production. Frontiers in Human Neuroscience, 18. 10.3389/fnhum.2024.1383714

Manes, J. L., Parkinson, A. L., Larson, C. R., Greenlee, J. D., Eickhoff, S. B., Corcos, D. M., & Robin, D. A. (2014). Connectivity of the subthalamic nucleus and globus pallidus pars interna to regions within the speech network: A meta-analytic connectivity study. Human Brain Mapping, 35(7), Article 7. 10.1002/hbm.22417

Müller, F., Nienstedt, J. C., Buhmann, C., Hidding, U., Gulberti, A., Pötter-Nerger, M., & Pflug, C. (2025). Effect of subthalamic and nigral deep brain stimulation on speech and voice in Parkinson’s patients. Journal of Neural Transmission, 132(3), 419–429. 10.1007/s00702-024-02860-5

Narayana, S., Jacks, A., Robin, D. A., Poizner, H., Zhang, W., Franklin, C., Liotti, M., Vogel, D., & Fox, P. T. (2009). A noninvasive imaging approach to understanding speech changes following deep brain stimulation in Parkinson’s disease. American Journal of Speech-Language Pathology, 18(2), 146–161. 10.1044/1058-0360(2008/08-0004)

Nozari, N. (2022). The Oxford Handbook of The Mental Lexicon. Oxford University Press.

Prasad, A. A., & Wallén-Mackenzie, Å. (2024). Architecture of the subthalamic nucleus. Communications Biology, 7(1), Article 1. 10.1038/s42003-023-05691-4

Remple, M. S., Bradenham, C. H., Kao, C. C., Charles, P. D., Neimat, J. S., & Konrad, P. E. (2011). Subthalamic Nucleus Neuronal Firing Rate Increases with Parkinson’s Disease Progression. Movement Disorders : Official Journal of the Movement Disorder Society, 26(9), Article 9. 10.1002/mds.23708

Rogalsky, C., Poppa, T., Chen, K.-H., Anderson, S. W., Damasio, H., Love, T., & Hickok, G. (2015). Speech repetition as a window on the neurobiology of auditory-motor integration for speech: A voxel-based lesion symptom mapping study. Neuropsychologia, 71, 18–27. 10.1016/j.neuropsychologia.2015.03.012

Rohl, A., Gutierrez, S., Johari, K., Greenlee, J., Tjaden, K., & Roberts, A. (2022). Chapter 7—Speech dysfunction, cognition, and Parkinson’s disease. In N. S. Narayanan & R. L. Albin (Eds.), Progress in Brain Research (Vol. 269, pp. 153–173). Elsevier. 10.1016/bs.pbr.2022.01.017

Roussel, P., Godais, G. L., Bocquelet, F., Palma, M., Hongjie, J., Zhang, S., Kahane, P., Chabardès, S., & Yvert, B. (2019). Acoustic contamination of electrophysiological brain signals during speech production and sound perception (p. 722207). bioRxiv. 10.1101/722207

Santens, P., De Letter, M., Van Borsel, J., De Reuck, J., & Caemaert, J. (2003). Lateralized effects of subthalamic nucleus stimulation on different aspects of speech in Parkinson’s disease. Brain and Language, 87(2), Article 2. 10.1016/S0093-934X(03)00142-1

Seidler, R. D., Noll, D. C., & Thiers, G. (2004). Feedforward and feedback processes in motor control. NeuroImage, 22(4), 1775–1783. 10.1016/j.neuroimage.2004.05.003

Shahid, S., Walker, J., Lyons, G. M., Byrne, C. A., & Nene, A. V. (2005). Application of higher order statistics techniques to EMG signals to characterize the motor unit action potential. IEEE Transactions on Biomedical Engineering, 52(7), 1195–1209. 10.1109/TBME.2005.847525

Simmonds, A. J., Leech, R., Collins, C., Redjep, O., & Wise, R. J. S. (2014). Sensory-Motor Integration during Speech Production Localizes to Both Left and Right Plana Temporale. The Journal of Neuroscience, 34(39), 12963–12972. 10.1523/JNEUROSCI.0336-14.2014

Tabari, F., Berger, J. I., Flouty, O., Copeland, B., Greenlee, J. D., & Johari, K. (2024). Speech, voice, and language outcomes following deep brain stimulation: A systematic review. PloS One, 19(5), e0302739. 10.1371/journal.pone.0302739

Tankus, A., & Fried, I. (2019a). Degradation of Neuronal Encoding of Speech in the Subthalamic Nucleus in Parkinson’s Disease. Neurosurgery, 84(2), Article 2. 10.1093/neuros/nyy027

Tankus, A., & Fried, I. (2019b). Degradation of Neuronal Encoding of Speech in the Subthalamic Nucleus in Parkinson’s Disease. Neurosurgery, 84(2), 378. 10.1093/neuros/nyy027

Temel, Y., Blokland, A., Steinbusch, H. W. M., & Visser-Vandewalle, V. (2005). The functional role of the subthalamic nucleus in cognitive and limbic circuits. Progress in Neurobiology, 76(6), 393–413. 10.1016/j.pneurobio.2005.09.005

Tjaden, K. (2008). Speech and Swallowing in Parkinson’s Disease. Topics in Geriatric Rehabilitation, 24(2), 115–126. 10.1097/01.TGR.0000318899.87690.44

Truccolo, W., Eden, U. T., Fellows, M. R., Donoghue, J. P., & Brown, E. N. (2005). A Point Process Framework for Relating Neural Spiking Activity to Spiking History, Neural Ensemble, and Extrinsic Covariate Effects. Journal of Neurophysiology, 93(2), 1074–1089. 10.1152/jn.00697.2004

Vissani, M., Bush, A., Lipski, W. J., Bullock, L., Fischer, P., Neudorfer, C., Holt, L. L., Fiez, J. A., Turner, R. S., & Richardson, R. M. (2025). Spike-phase coupling of subthalamic neurons to posterior perisylvian cortex predicts speech sound accuracy. Nature Communications, 16(1), 3357. 10.1038/s41467-025-58781-8

Wang, E. Q., Metman, L. V., Bakay, R. A. E., Arzbaecher, J., Bernard, B., & Corcos, D. M. (2006). Hemisphere-Specific Effects of Subthalamic Nucleus Deep Brain Stimulation on Speaking Rate and Articulatory Accuracy of Syllable Repetitions in Parkinson’s Disease. Journal of Medical Speech-Language Pathology, 14(4), Article 4.

Watson, P., & Montgomery, E. B. (2006). The relationship of neuronal activity within the sensori-motor region of the subthalamic nucleus to speech. Brain and Language, 97(2), Article 2. 10.1016/j.bandl.2005.11.004

Weismer, G. (2023). Oromotor Nonverbal Performance and Speech Motor Control: Theory and Review of Empirical Evidence. Brain Sciences, 13(5), 768. 10.3390/brainsci13050768

